# Coupling Day Length Data and Genomic Prediction tools for Predicting Time-Related Traits under Complex Scenarios

**DOI:** 10.1101/703488

**Authors:** Diego Jarquin, Hiromi Kajiya-Kanegae, Chen Taishen, Shiori Yabe, Reyna Persa, Jianming Yu, Hiroshi Nakagawa, Masanori Yamasaki, Hiroyoshi Iwata

**Affiliations:** Department of Agronomy and Horticulture, University of Nebraska – Lincoln, Lincoln NE, 68583, USA; Graduate School of Agricultural and Life Sciences, The University of Tokyo, Bunkyo, Tokyo 113-8657, Japan; Institute of Crop Sciences, National Agriculture and Food Research Organization (NARO), Tsukuba, Ibaraki 305-8518, Japan; Department of Agronomy, Iowa State University, Ames, USA; Food Resources Education and Research Center, Graduate School of Agricultural Science, Kobe University, Kasai, Hyogo 675-2103, Japan; Institute for Agro-Environmental Sciences, National Agriculture and Food Research Organization (NARO), Tsukuba Ibaraki 305-8604

**Author notes:** Corresponding authors: University of Nebraska – Lincoln, Lincoln NE, 68583, USA. (DJ). Graduate School of Agricultural and Life Sciences, The University of Tokyo, Bunkyo, Tokyo 113-8657, Japan., (HI).

## Abstract

Genomic selection (GS) has proven to be an efficient tool for predicting crop-rank performance of untested genotypes; however, when the traits have intermediate optima (phenology stages) this implementation might not be the most convenient. GS might deliver high rank correlations but incurring in serious bias. Days to heading (DTH) is a crucial development stage in rice for regional adaptability with significant impact in yield potential. The objectives of this research consisted in the development of a method that accurately predicts time related traits like DTH in unobserved environments. For this, we developed an implementation that incorporate day length information (DL) in the prediction process for two relevant scenarios: CV0, predicting tested genotypes in unobserved environments (C method); and CV00, predicting untested genotypes in unobserved environments (CB method). The use of DL has advantages over weather data since it can be determined in advance just by knowing the location and planting date. The proposed methods showed that DL information significantly helps to improve the predictive ability of DTH in unobserved environments. Under CV0, the C method returned an root-mean-square-error (RMSE) of 3.9 days, a Pearson Correlation (PC) of 0.98 and the differences between the predicted and observed environmental means (EMD) ranged between −4.95 and 4.67 days. For CV00, the CB method returned an RMSE of 7.3 days, a PC of 0.93 and the EMD ranged between −6.4 and 4.1 days while the conventional GS implementation produced an RMSE of 18.1 days, a PC of 0.41 and the EMD ranged between −31.5 and 28.7 days.

## Introduction

To satisfy the daily demand of crop consumption for a growing population, it is necessary to improve the production systems to allow a sustainable increase of the yield potential and nutritional value of the new and up-coming varieties^1^. In plant breeding there are different tools and methodologies to increase the rate of genetic gain^2^. Traditional plant breeding methods use phenotypic and/or pedigree information to perform selection of experimental lines for developing superior breeding lines. However, phenotypic selection may not be permissible in many situations due to the high phenotyping costs; also, pedigree selection may not be accurate since it assumes a constant rate of recombination (50% of the genetic material coming from mother and the remaining 50% coming from father) and this might not be a feasible premise. In this regard, the use of molecular markers can help to detect the deviations from the expected genetic material of the parents and these can be used to determine which parental haplotypes were inherited.

Genomic selection^3^ (GS) is an emerging tool that allows screening genotypes from a very large population without having to observe them in fields^4,5^. This method only requires phenotypic and genomic information for calibrating models, then other genotyped candidates are selected based on the predicted values obtained via their marker profiles^6^. In this context, the implementation of genomic tools and resources translates our understanding of the relationship between genotype and phenotype into predicted genetic merits for direct selection. This is highly desirable dealing with those traits that are controlled by a large number of genes with small individual effects, also known as complex traits^5,7^. GS has shown to be an effective tool for plant and animal breeding applications, especially dealing with complex traits^8^. The successfulness of GS predicting phenotypes from genotypic information has been reported in various crop species^9^.

In GS, the genomic estimated breeding values (GEBVs) of complex traits of untested genotypes are computed based on genome-wide marker information. The GEBVs can be considered as deviations around a fixed mean (e.g., zero). Once the GEBVs are obtained, the breeders may conduct selections based on the ranked values by selecting those individuals in the tails. Depending on the trait, the individuals are selected either on the upper or lower tails of the ranked GEBVs. For example, the upper side is selected for increasing genetic gains of traits like grain yield, stress resistance, or the content of essential nutrients. On the other hand, genotypes in the lower tail are selected for reducing lodging, shattering, the content of anti-nutrients, etc. In these cases, the EBVs provide sufficient information when the correlation between the EBVs and the direct genetic values (or phenotypic values) is high. However, the rank-based selection criteria might not be the most accurate when the target trait has an intermediate optima. This is the case of those traits related to phenological stages of crop plants. For example, Onogi *et al*^10^., the prediction of heading date of untested genotypes within and across environments by integrating genomic prediction with crop/ecophysiological modelling. These authors predicted DTH for untested genotypes within environments and for untested genotypes in unobserved environments among other cross-validations schemes. The corresponding root-mean-square-error (RMSE) predicting untested genotypes in unobserved environments was almost twice than for the within environments prediction schemes. Moreover, in both cases high correlations between predicted and observed values were obtained. For the within environments the correlation was of 0.96 while for predicting untested genotypes in unobserved environments it was of 0.87. These results highlight the importance of improving the accuracy of predicting untested genotypes in unobserved environments despite the fact of obtaining high correlations.

In rice, days to heading (DTH; typically defined as the number of days after planting when more than 50% of individuals show the heading stage) is one of the critical traits of ideotypes with an increased yield potential. Hence it is important to optimize DTH for high yield potential in different environments^11^. Usually multi environment trials (METs) are performed for developing stable cultivars across a wide range of environmental conditions, and the breeders may be interested in estimating DTH for a large set of environments instead of a single one. However, when the GEBVs are computed for an unobserved environment these might contain a large bias if the phenotypic information of genotypes in the training set do not resemble the target environment. The GEBVs of one environment may not be valid for another environment due to variations in the environmental *stimuli* causing inconsistencies in the response patters as result of the interaction between genotypes to environments (G×E interaction). These differences remain also for same genotypes observed in same locations but tested at different times (planting date, years). As consequence of the aforementioned issues, the problem of correctly estimating DTH for a set of untested genotypes in a wide range of unobserved environments becomes harder to solve than for non-time related traits. As pointed before, the advantage of GS techniques over conventional phenotypic methods is its ability for performing accurate rank predictions of untested genotypes based exclusively on their marker profiles without having to observe these in fields. Thus, a method capable of delivering highly accurate predictions of untested genotypes in unobserved environments without incurring in a large bias is desirable.

As was pointed before, DTH in rice is important for regional adaptability and has usually great impact on yield potential^11^, genetic systems controlling it have been studied for a long time by integrating Mendelian genetics and genomics^12^. The potential of GS in DTH has been evaluated in rice^13,14^ and sorghum^15^. Onogi *et al*.^10^ focused on the genomic prediction of DTH in multiple environments and demonstrated the potential of a novel method that integrates GS with a crop growth model. Li *et al.^15^* also focused on the genomic prediction of DTH in multiple environments and demonstrated the potential of models combining GS with joint regression analysis on photothermal time. To our knowledge, no studies combining DL at the day when genotypes reach DTH with GS tools have been done before. The objectives of this research consisted in the development of a method that accurately predicts time related traits like DTH in unobserved environments by leveraging information from day length and molecular markers.

Two cross-validation schemes of interest for designing and planning experiments were considered for evaluating the performance of the proposed implementation for predicting DTH in unobserved environments: CV0, predicting already tested genotypes in unobserved environments (targeting environments); and CV00, predicting untested genotypes in unobserved environments (targeting genotypes and environments). For assessing the proposed implementation the root-mean-square-error (RMSE), the Pearson correlation (PC) and the difference between predicted and observed environmental means (EMD) values were computed.

In this study, we propose a novel methods to predict DTH in a precise and accurate way (i.e., small bias and high correlation between predicted and observed values) of tested (C method) and untested (CB method) genotypes in unobserved environments.

Under the hardest cross-validation scheme (i.e., CV00), the results of the introduced CB method are compared with those obtained using the conventional GS implementation (G-BLUP). For studying the realistic scenario of predicting already tested genotypes in unobserved environments, the cross validation scheme CV0 was considered. These results are used for assessing the levels of predictability that can be reached when phenotypic information of a genotype is available in other environments. Here the environments are considered “target” in a strict sense because these can be selected even before conducting field experiments using the theoretical DL values of locations for a given planting date.

## Results

### Descriptive statistics

The box-plot of DTH is depicted in Figure S1. The environments were ordered based on the environmental medians. In general, those environments with late planting date appear first. The medians ranged between 59 (“Fukuyama 2010 Late”) and 116 (“Akita 2015”) days. The SDs of the environments varied between 5.07 and 16.83 days. Also, the smallest and the largest DTH values were 42 and 148 days and these were observed in “Tsukubamirai 2016 Late” and “Akita 2015” environments, respectively.

### Evaluating the bias and accuracy of the methods

Table 1 contains the observed and the predicted environmental means (E-mean) for two different cross-validation schemes (CV00 and CV0) as well as the corresponding RMSE and the PC between predicted and observed values. For the CV00 scheme, the CB and GS methods were implemented. The objective of using this cross-validation scheme was to assess the model performance of the proposed method with respect to the conventional GS implementation for predicting untested genotypes in unobserved environments. Since no phenotypic information is available for connecting yet to be predicted genotypes in unobserved environments with genotypes in training sets, the CV00 scheme is considered the hardest and the most interesting prediction problem. Across environments, the conventional GS method returned an RMSE of 18.1 days and a PC of 0.41; within environments, the average RMSE and PC were of 17.5 days and 0.69. The EMD ranged between −31.5 and 28.7 days. On the other hand, across environments the proposed CB method returned an RMSE of 7.3 days and a PC of 0.93; within environments, the average RMSE and PC were of 7.2 days and 0.86. The EMD ranged between −6.4 and 4.1 days.

**Table 1.** Environmental means, Root Means Squared Error (RMSE) and the Pearson correlation between predicted and observed days to heading (DTH) for 112 rice genotypes tested in 51 environments in Japan. Two cross-validation schemes were implemented for mimicking realistic prediction scenarios: CV0 represents the scenario of predicting tested genotypes in unobserved environments; CV00 considers the prediction of untested genotypes in unobserved environments. For CV0, the C method was used combining phenotypic and day length information (DL) of tested genotypes (one at a time) in other environments. For CV00, two methods were considered: (*i*) the conventional GS implementation; and (*ii*) the CB method, which combines phenotypic and DL information from the tested genotypes (training set) and genomic BLUP values (*ĝ*) of the untested genotypes. In both cases, the prediction procedure was conducted by leaving one genotype out across environments and deleting any phenotypic information from the target environment as well.

|  | CV00 |  |  |  |  |  |  | CV0 |  |  |
| --- | --- | --- | --- | --- | --- | --- | --- | --- | --- | --- |
|  | Mean |  |  | RMSE |  | Corr |  | Mean | RMSE | Corr |
|  | Real | CB | GS | CB | GS | CB | GS | C | C | C |
| Tsukubamirai 2004 Late | 79.4 | 76.7 | 85.9 | 7.0 | 12.6 | 0.86 | 0.70 | 76.3 | 4.6 | 0.99 |
| Tsukubamirai 2004 Early | 87.5 | 93.2 | 84.1 | 10.0 | 12.9 | 0.84 | 0.55 | 92.2 | 6.4 | 0.97 |
| Tsukubamirai 2005 Early | 98.5 | 102.1 | 87.1 | 6.9 | 14.0 | 0.87 | 0.75 | 101.0 | 3.5 | 0.98 |
| Tsukubamirai 2005 Late | 76.7 | 74.8 | 84.8 | 5.7 | 12.4 | 0.86 | 0.68 | 73.7 | 3.4 | 0.99 |
| Kasai 2006 | 87.4 | 83.8 | 88.3 | 7.9 | 8.8 | 0.85 | 0.75 | 82.8 | 6.8 | 0.93 |
| Fukuyama 2006 | 80.6 | 83.0 | 82.5 | 5.6 | 13.4 | 0.87 | 0.29 | 82.0 | 2.0 | 0.99 |
| Fukuyama 2007 | 84.1 | 83.4 | 86.6 | 5.0 | 6.1 | 0.87 | 0.84 | 82.5 | 2.9 | 0.98 |
| Tsukuba 2008 | 104.5 | 107.7 | 87.2 | 9.1 | 19.2 | 0.87 | 0.85 | 107.3 | 4.0 | 0.98 |
| Kasai1st 2008 | 102.2 | 99.9 | 88.7 | 9.0 | 17.8 | 0.89 | 0.71 | 99.2 | 4.3 | 0.99 |
| Fukuyama 2008 | 77.8 | 82.2 | 85.6 | 7.8 | 11.1 | 0.86 | 0.77 | 81.3 | 4.5 | 0.98 |
| Tsukuba 2009 | 103.9 | 107.7 | 87.1 | 8.6 | 19.0 | 0.89 | 0.81 | 108.0 | 5.2 | 0.98 |
| Kasai1st 2009 | 101.9 | 100.6 | 89.4 | 7.6 | 15.1 | 0.89 | 0.83 | 100.0 | 2.7 | 0.99 |
| Fukuyama 2009 | 78.7 | 78.7 | 89.6 | 5.2 | 13.2 | 0.86 | 0.83 | 77.8 | 1.5 | 0.99 |
| Tsukuba 2010 Late | 83.5 | 89.1 | 88.9 | 8.4 | 11.1 | 0.86 | 0.66 | 88.4 | 5.9 | 0.97 |
| Kasai1st 2010 | 100.6 | 100.7 | 89.5 | 7.1 | 14.3 | 0.88 | 0.77 | 100.1 | 1.9 | 0.99 |
| Fukuyama 2010 | 76.2 | 79.5 | 88.2 | 5.9 | 15.3 | 0.86 | 0.50 | 78.1 | 2.6 | 0.99 |
| Fukuyama 2010 Late | 59.0 | 65.4 | 87.7 | 9.2 | 29.7 | 0.84 | 0.82 | 62.5 | 4.2 | 0.96 |
| Tsukuba 2011 Early | 105.9 | 109.9 | 85.5 | 9.9 | 23.9 | 0.87 | 0.66 | 109.9 | 5.3 | 0.98 |
| Tsukuba 2011 Late | 82.8 | 80.8 | 90.0 | 5.9 | 11.9 | 0.86 | 0.78 | 80.4 | 4.1 | 0.95 |
| Tsukuba 2011 Middle | 94.7 | 95.7 | 90.9 | 6.5 | 11.1 | 0.88 | 0.67 | 95.4 | 3.0 | 0.98 |
| Kasai 2011 | 101.4 | 100.6 | 89.3 | 7.8 | 15.7 | 0.89 | 0.76 | 100.0 | 2.4 | 0.99 |
| Fukuyama 2011 | 75.7 | 79.4 | 91.1 | 6.4 | 16.9 | 0.87 | 0.83 | 78.6 | 3.1 | 0.99 |
| Fukuyama 2011 Late | 66.4 | 72.7 | 97.9 | 8.9 | 32.4 | 0.68 | 0.58 | 66.2 | 2.9 | 0.95 |
| Tsukuba 2012 Early | 97.1 | 96.0 | 86.9 | 5.7 | 12.4 | 0.81 | 0.74 | 101.1 | 5.8 | 0.97 |
| Tsukuba 2012 Late | 84.3 | 81.6 | 88.2 | 6.1 | 7.5 | 0.86 | 0.80 | 81.0 | 4.3 | 0.96 |
| Tsukuba 2012 Middle | 92.6 | 95.4 | 88.4 | 6.3 | 8.3 | 0.87 | 0.82 | 94.8 | 3.7 | 0.97 |
| Kasai 2012 | 103.8 | 102.7 | 86.4 | 7.9 | 21.3 | 0.89 | 0.60 | 102.1 | 2.7 | 0.99 |
| Fukuyama 2012 | 75.2 | 78.8 | 89.9 | 6.2 | 16.3 | 0.87 | 0.83 | 78.0 | 3.1 | 0.99 |
| Fukuyama 2012 Late | 71.5 | 77.0 | 103.1 | 7.5 | 32.4 | 0.70 | 0.66 | 68.6 | 3.4 | 0.98 |
| Fukuoka 2012 | 79.6 | 76.0 | 89.1 | 6.7 | 13.0 | 0.84 | 0.69 | 75.4 | 5.2 | 0.96 |
| Akita 2012 | 113.1 | 115.9 | 90.1 | 8.3 | 26.1 | 0.86 | 0.54 | 114.5 | 2.8 | 0.98 |
| Kasai 2013 | 104.8 | 104.1 | 87.3 | 8.0 | 20.1 | 0.90 | 0.80 | 103.5 | 3.2 | 0.98 |
| Fukuyama 2013 | 74.9 | 80.1 | 89.4 | 7.4 | 16.3 | 0.87 | 0.82 | 79.3 | 4.6 | 0.99 |
| Fukuyama 2013 Late | 62.1 | 64.8 | 85.7 | 6.3 | 27.1 | 0.83 | 0.36 | 64.1 | 3.6 | 0.93 |
| Fukuoka 2013 | 77.4 | 77.3 | 93.3 | 5.7 | 17.5 | 0.86 | 0.77 | 76.7 | 1.9 | 0.99 |
| Akita 2013 | 116.6 | 115.8 | 91.1 | 8.5 | 28.1 | 0.88 | 0.66 | 115.1 | 2.4 | 0.99 |
| Kasai 2014 | 102.4 | 105.1 | 88.9 | 8.4 | 18.0 | 0.90 | 0.66 | 104.4 | 2.8 | 0.99 |
| Fukuyama 2014 | 79.6 | 82.1 | 87.9 | 5.9 | 12.1 | 0.88 | 0.68 | 81.3 | 2.1 | 0.99 |
| Fukuyama 2014 Late | 67.6 | 64.5 | 88.1 | 6.7 | 21.4 | 0.83 | 0.79 | 62.9 | 5.1 | 0.98 |
| Fukuoka 2014 | 81.8 | 77.7 | 80.4 | 6.7 | 11.5 | 0.87 | 0.45 | 77.1 | 5.4 | 0.97 |
| Akita 2014 | 115.2 | 116.2 | 89.6 | 9.2 | 28.5 | 0.88 | 0.68 | 115.6 | 2.2 | 0.99 |
| Fukuyama 2015 | 76.3 | 72.6 | 88.1 | 6.5 | 14.4 | 0.86 | 0.63 | 71.9 | 4.6 | 0.99 |
| Akita 2015 | 118.1 | 119.8 | 89.4 | 7.7 | 29.9 | 0.81 | 0.76 | 117.6 | 2.5 | 0.99 |
| Kasai 2016 | 102.3 | 101.2 | 85.6 | 7.2 | 20.4 | 0.90 | 0.61 | 100.7 | 3.1 | 0.98 |
| Fukuyama 2016 | 79.1 | 81.1 | 85.2 | 6.0 | 8.9 | 0.86 | 0.82 | 80.8 | 2.4 | 0.99 |
| Fukuyama 2016 Late | 67.5 | 65.2 | 89.6 | 6.8 | 26.1 | 0.79 | 0.63 | 64.6 | 4.5 | 0.95 |
| Tsukubamirai 2016 Late | 73.6 | 77.9 | 84.6 | 7.0 | 13.6 | 0.90 | 0.80 | 77.2 | 4.9 | 0.98 |
| Kasai 2017 | 108.6 | 106.2 | 85.7 | 8.3 | 26.3 | 0.89 | 0.56 | 105.5 | 4.3 | 0.98 |
| Fukuyama 2017 | 77.4 | 80.2 | 87.7 | 6.3 | 16.3 | 0.88 | 0.42 | 79.6 | 2.7 | 0.99 |
| Fukuyama 2017 Late | 68.4 | 67.7 | 88.8 | 6.0 | 22.9 | 0.82 | 0.69 | 67.0 | 3.0 | 0.97 |
| Tsukubamirai 2017 Late | 69.5 | 73.1 | 86.9 | 6.8 | 19.8 | 0.87 | 0.65 | 72.2 | 4.3 | 0.98 |
| Mean |  |  |  | 7.2 | 17.5 | 0.86 | 0.69 |  | 3.7 | 0.98 |

The C method was implemented under the CV0 scheme and their results were used for assessing the predictive ability of predicting an already tested genotype in an untested environments considering only the phenotypic and DL information of the same genotype. Across environments, this method returned an RMSE of 3.9 days and a PC of 0.98; within environments, the average RMSE and PC were of 3.7 days and 0.98. The EMD ranged between - 4.95 and 4.67 days.

The results of CB and GS methods are directly comparable between them (CV00 scheme). Figure 1 depicts the scatter plot between predicted and observed DTH values for all genotypes across environments. The top panel contains the results obtained with the proposed CB method, while the bottom panel shows the results derived from the conventional GS implementation. The blue and black lines correspond to the fitted and the on target predictions with respect to the observed DTH values. In general, across environments (Figure 1) and within environments (Table 1) the results obtained with the CB method showed a better fit than those derived from the GS implementation.

**1. Figure 1.**
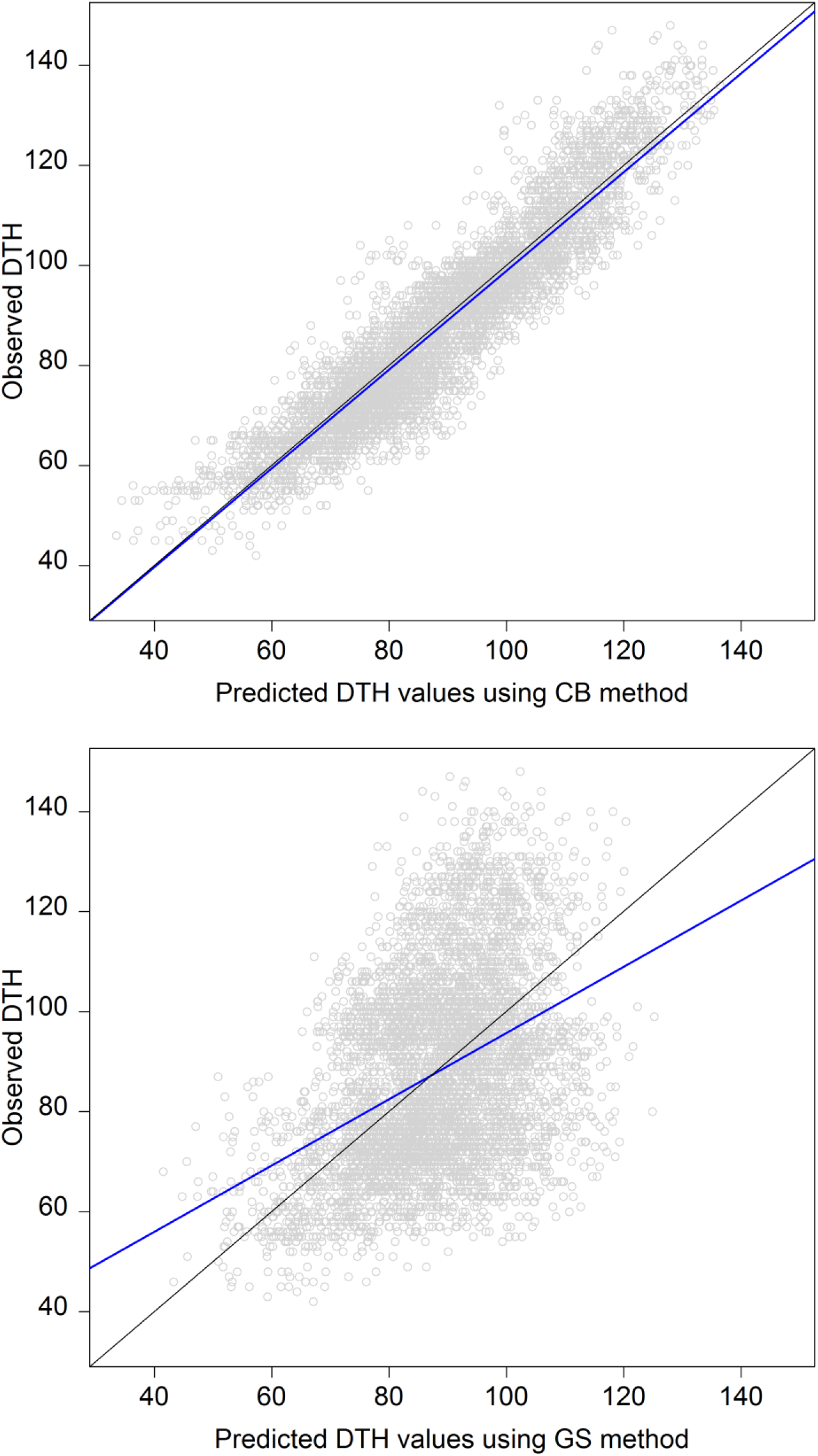
Scatter plot of predicted (*x-axis*) and observed (*y-axis*) DTH values under CB (top) and GS (bottom) methods for the CV00 scheme (predicting untested genotypes in unobserved environments) for a rice data set comprising 112 genotypes tested in 51 environment in Japan between 2005 and 2017. The CB method combines the predicted environmental means (E-mean) and the genomic BLUPs (*ĝ*) derived from the conventional GS implementation while the GS implementation considers a common mean across environments plus the genomic BLUPs. The blue line represents the fitted line between predicted and observed DTH values, and the black line show the exact match (on target) of the predicted values on the observed values.

The advantages of CB method for predicting DTH are clearly appreciated considering the predicted environmental means (E-means). Figure 2 displays the predicted (*x-axis*) and observed (*y-axis*) E-means under the CB (orange) and GS (blue) methods. As expected, when no information from the target environment was available the conventional GS implementation predicted the DTH values to be centered on the overall mean (~88 days) of the environments observed in training set. On the other hand, the CB method consistently returned closer values to the diagonal black line, which represents the exact match.

**2. Figure 2.**
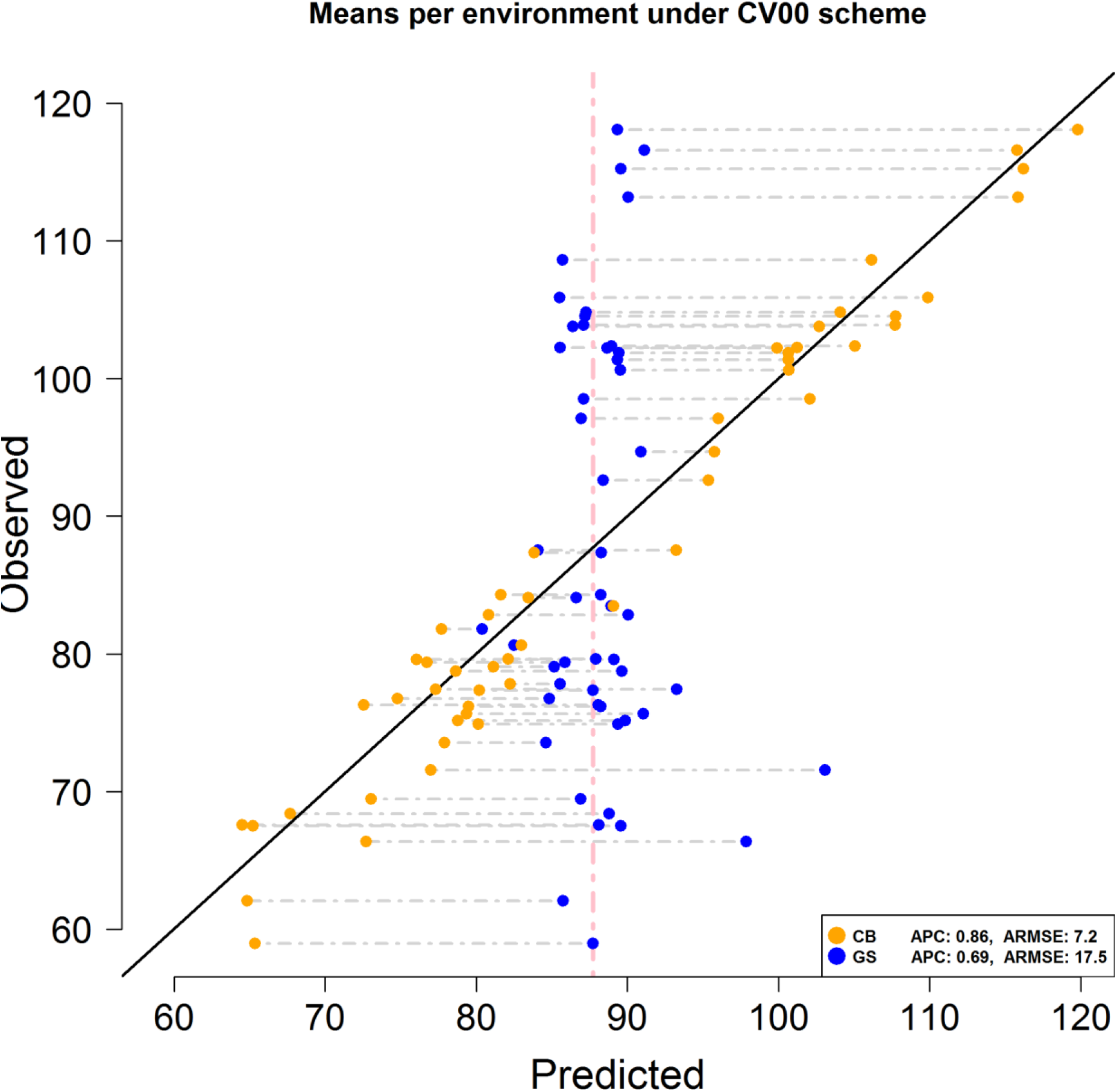
Scatter plot of predicted and observed environmental means (E-means) using the CB method (orange) and the conventional GS implementation (blue) for the CV00 scheme (predicting untested genotypes in unobserved environments) for a rice data set comprising 112 genotypes tested in 51 environment in Japan between 2005 and 2017. The vertical pink line indicates the position of the overall phenotypic mean across environments (88 days). The diagonal black line shows the exact match between predicted and observed means. ARMSE and ACP stand for the Average Root Mean Square Error and the Average Person Correlation between predicted and observed DTH values across environments. The RMSE and PC were first computed within environments and the obtained values were averaged.

## Discussion

The boxplot (Figure S1) showed a high heterogeneity among the environments in the study with those with “late” planting date exhibiting smaller phenotypic variability and shorter occurrence time for DTH compared with the “early” planting environments. Among other considerations, the conventional GS models assume homogeneity of variances between environments. However, the violation of this assumption may induce serious inconsistencies in the results. Especially when genotype-by-environment interaction parameters are not explicitly included or accounted in models.

The results from GS model provide an overview of the possible problems that we might incur predicting time related traits when no phenotypic information from the target environment is available (CV00). This cross-validation scheme represents among the four prediction scenarios that breeders face in field (CV2: predicting tested genotypes in observed environments; CV1: predicting untested genotypes in observed environments; CV0: predicting tested genotypes in unobserved environments; and CV00: predicting untested genotypes in unobserved environments) the most interesting, useful and challenging. Dealing with traits where the selection process is based in ranks rather than in the *per se* estimated values, the conventional implementation (GS) usually provides good results (moderate to high correlations) under the four schemes.

However, in traits related with important growth stages like DTH where the *per se* estimated values are crucial for designing, planning and establishing future experiments (i.e., CV0 and CV00 situations) the implementation of the conventional implementations (GS) might not be the best choice. When no phenotypic information of the target environment is available, the E-mean of the target environment is determined as the weighted average of those environments in the training set forcing it to be centered towards a general mean corresponding to the weighted mean of those environments in the training set. Hence, the final prediction is composed of this general weighted mean plus deviations corresponding to the estimated/predicted genomic effects. Since these deviations are conceptualized to be centered round zero, under CV0 and CV00 schemes the GS method would produce similar environmental means (i.e., centered on an overall mean determined by the environments in the calibration set).

The results from the C method showed that this implementation provides good estimations of DTH for the CV0 scheme. Across environments, the RMSE was of 3.9 days and for a PC of 0.98. These results were very similar to those obtained using the conventional GS models when complete information of the genotypes across environments was available (i.e., CV2 scheme). Under CV2, the SMSE and Pearson correlation across environment were of 3.8 days and 0.98, respectively (data not shown).

As mentioned before, it is expected to obtain similar environmental means using either CV0 or CV00 schemes under the conventional GS implementations. Under the CV00 scheme, the GS strategy returned a RMSE of 18.1 and a PC of 0.41 across environments. The EMD between predicted and observed values ranged between −31.5 and 28.7 days. On the other hand, the C method returned an EMD ranging between −4.95 and 4.67 days under the CV0 scheme. In this case, only the phenotypic information of the genotype to be predicted in an observed environment but tested in others was used. Since under the CV0 and CV00 schemes the conventional GS implementation would produce similar E-means when no information of the target environment is available then it is also expected that the proposed C method will produce better results than those derived from the GS implementation under the CV0 scheme.

As was pointed before, one of the main goals of this research was to develop a method able to deliver small RMSE between predicted and observed values when predicting DTH of untested genotypes in unobserved environments (CV00 scheme). Under this scenario the CB method produced encouraging results. The RMSE across environments was of 7.3 days and a PC of 0.86. A significant improvement was achieved compared with those results obtained using the conventional GS implementation (RMSE of 18.1 days and PC of 0.41). In this case, the CB method doubled the accuracy of the conventional GS method and reduced the SMSE by around 60%. However, the advantages of using this method under CV00 scheme can be appreciated better when comparing the predicted environmental means. Under the conventional GS implementation, the differences between predicted and observed environmental means ranged between −31.5 and 28.7 days while with the CB method these only ranged between −6.4 and 4.1 days. The CB method reduced the extremes (left and right) of the range obtained with the GS implementation in between 80 and 85%, respectively. The advantages of the CB method are evident in Figure 2 were their environmental means (orange dots) were always closer to the diagonal line that represents the exact match with true means compared with those means from the conventional GS implementation (blue dots).

At first, the implementation of the C and CB methods requires enough information for accurately characterize the genotype’s DL curves. For this, sets of genotypes tested in several environments are necessary. However, once a good characterization of the DL curves is accomplished potentially any untested genotypes may be accurately predicted in any unobserved environment. Thus, the obtained results should be accurate not only for the set of genotypes and environments analyzed in this study but also for others.

The methods here introduced (C and CB) showed that the use of DL information significantly improves the predictive ability of the conventional GS implementations with a significant reduction of the RSME between the predicted and observed DTH values. An advantage of the C method is that if phenotypic information of the same genotype is available for a set of environments, it is easy to estimate the expected DTH in an unobserved environment without needing molecular marker information or phenotypic information from other genotypes. The CB method improved the prediction of DTH of untested genotypes in unobserved environments by coupling the marker profiles of the untested genotypes with the phenotypic and DL information from other genotypes.

Both of the introduced methods require the day length (DL) information of the environments (training and testing) in their design. The convenience of these methods resides in the fact that the theoretical DL information can be obtained in advance before establishing and designing the experiments thus the day length data can be help breeders to schedule the crop season better. The proposed methods capture the genotype-by-environment interaction (G×E) component via the fitted curves thus the explicitly inclusion of this term is not needed. The conventional implementation (GS) returns the same estimated genomic effect across environments for a particular genotype when the G×E component is absent. The curves precisely capture the sensitivity experienced by genotypes under different environments driving specific responses to each environment. Thus, although the CB method is composed of the genomic component computed without explicitly considering the G×E term the final prediction accounts for this interaction term via the curves. In addition, the formal modeling of heterogeneity of variances is not necessary because these patterns are also accounted with the current method via the curves.

## Materials and Methods

### Phenotypic data

The phenotypic data set comprises the seeding and heading date of 112 Japanese rice cultivars with different origin from different regions and these correspond to improved lines and landraces (Table S1). These genotypes were tested in 51 locations (North to South of Japan) from 2004 to 2017 (Table S2). This multi-environmental trials (METs) experiment was conducted for investigating heading date related genes. The materials had polymorphisms known heading date related genes (e.g., Hd1, Hd2, Hd5 and Hd6).

The number of DTH was calculated as the difference in days between the heading date and the seeding date. The heading date was determined when more than 50% of the individuals in the plot reached the heading stage.

### Marker genotype data

Details procedures of DNA extraction and library construction for whole genome shotgun sequence are described by Yabe *et al*^17^. For each variety, total DNA was extracted using the CTAB method^18^. The library was constructed using the Illumina TruSeq DNA Sample preparation kit according to manufacturer’s instructions (Illumina, Inc., San Diego, CA, USA). Next-generation sequencing data was obtained using the Illumina HiSeq 2000, HiSeq 4000 and HiseqX systems via paired-end sequencing (Illumina, Inc., San Diego, CA, USA). The sequence data have been deposited in the DNA Data Bank of Japan (DDBJ) Sequence Read Archive, under Submission DRA008071. Data sets deposited in the DDBJ Sequence Read Archive (SRA106223, ERA358140, DRA000158, DRA000307, DRA000897, DRA000927, DRA007273, DRA007256) were reanalyzed. Adapters and low-quality bases were removed from paired reads using the Trimmomatic v0.36 program^19^. The preprocessed reads were mapped to the Os-Nipponbare-Reference-IRGSP-1.0^20^ by using the bwa-0.7.12 mem algorithm with the default options^21^. The mean depth of the reads against the Os-Nipponbare-Reference-IRGSP-1.0 ranged from 7.4x to 72.4x with average of 16.9x.

SNP calling was based on alignment using the Genome Analysis Toolkit^22^ (GATK, 3.7-0-gcfedb67) and Picard package V2.5.0 (http://broadinstitute.github.io/picard). The mapped reads were realigned using RealignerTargetCreator and indelRealigner of GATK software. SNPs and InDel were called at the population level using the UnifiedGenotyper of GATK with the - glm BOTH option. Heterozygous loci were treated as missing, owing to the assumption that almost all markers were well fixed in rice cultivars after several generations of self-pollination. We used only biallelic non-missing sites over all cultivars with a minor allele frequency (MAF) ≥ 0.025. Initially, 629,602 SNP markers were considered in the study. After applying discarding those with more than 50% of missing values 408,372 SNPs remaining in the study for analysis.

### Day length data

Although there are several definitions for day length (DL) according to the Smithsonian Institution^23^ roughly speaking it can be defined as the amount of time in a day with sunlight. The theoretical DL values were computed by the default method^16^ included in the geosphere (v1.5-5) R package^24^. DL information can be known in advance just by knowing the location and date and it is not affected by the weather conditions.

### Cross-Validation Schemes

Two cross-validation schemes were considered in the study and these mimic different situations that breeders might face in fields. Figure S2 shows the graphical representation of CV0 (predicting tested genotypes in unobserved environments) and CV00 (predicting untested genotypes in unobserved environments). For CV0, the cross-validation is conducted by deleting the phenotypic information for all genotypes at each environment (one at a time) and using the remaining environments as training set. Using the information from Figure S2 (top panel), this procedure is sequentially repeated six times (one for each environment).

For implementing the CV00 scheme, for each environment in the testing set all the phenotypic information is deleted as in the previous cross-validation scenario; however, also sequentially the phenotypic information from each one of the genotypes in the testing environment is deleted from all environments (one at a time). In the bottom panel of Figure S2, we have that the goal was to predict genotype G3 in environment E4. For this environment, the same procedure is repeated for each one of the genotypes (G1-G5). Such that with six environments and five genotypes we have that this procedure is implemented 30 (5×6) times.

### GBLUP model

The Genomic Best Linear Unbiased Predictor model (Genomic BLUP or GBLUP) is a main effects model used in GS applications^25,8^. It attempts to explain the DTH value of the *i*^th^ genotype (*i*=1, 2,…, *I*) observed in the *j*^th^ environment (*j*=1, 2,…, *J*) as the sum of a common mean, random environmental and genomic effects, and an error term and it is represented as follows:

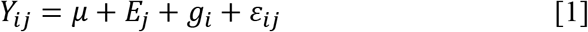

where *μ* is a common effect to all genotypes across all environments; *E_j_* are IID (independent and identically distributed) outcomes from a 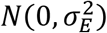 representing the effect of the *j*^th^ environment; 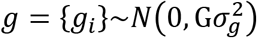 is the vector of genomic effects, with 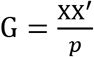 as the genomic relationship matrix whose entries describe the genetic similarities between pairs of individuals, X is the standardized (by columns) matrix and *p* is the number of molecular markers; *ε_ij_* are IID outcomes from a 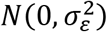 representing measure errors; and 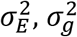 and 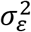 are the corresponding associated variance components of the described random effects.

Usually, when METs are analyzed the environmental component explains the larger proportion of the phenotypic variability across environments^26,27^. Hence, from [1] is concluded that the predicted value of the *i*^th^ genotype in the *j*^th^ environment would strongly depend on the environmental mean, which is composed by a common mean across environments plus an environmental deviation that corresponds to the effect of *j*^th^ environment.

Conceptually, the common mean effect can be viewed as the average performance of all genotypes across environments. The environmental deviation corresponds to the mean of the specific affectations of the current environmental *stimuli* in the set of the tested genotypes beyond the common mean. Thus, these environmental effects can be considered deviations around the common mean. Hence, by construction the contribution of the genomic component to the idealized phenotype is much smaller in comparison with the overall mean and the environmental effects.

If phenotypic information (DTH) of the target environment is available, good approximations of the environmental effects can be computed. Otherwise, the environmental effects are estimated using the phenotypic information of the genotypes observed in other environments. Thus, the environmental component is estimated as the weighted average of the environmental effects in the training set. Since these random effects are conceptualized to be centered on zero, the weighted average will be close to zero. Hence, the predicted DTH value will be compose mainly of a common mean and a small genomic effect shrinking the predicted values towards the common mean. Although this method usually delivers high correlations between predicted and observed values, it also might induce a large bias (RMSE) negatively affecting the accuracy of the prediction of time related traits.

For alleviate this, the two introduced methods (C and CB) incorporate DL information in the prediction process. The idea for using DL information in the prediction process was motivated when examining the relationship between the Cumulative Day Length (CDL) and DTH of all genotypes across environments (Figure S3). A simple linear regression of DTH on CDL returned an R-squared value of 0.99 exhibiting a strong relationship between these two variables. From there we can conclude that if CDL values where to be known then accurate estimations of DTH may be obtained for any genotype in any environment. However, in order to compute the CDL the exact number of days between planting date and the heading occurrence (DTH) should be known at first complicating the estimation process.

Since each genotype was observed in a considerable number of environments (48-50) these might show response patterns between DTH and DL when analyzing one at a time. In this study, we observed that across environments the DL values at the DTH occurrence showed strong relationship response patterns with DTH. For illustrating the conceptualization of this idea we use as example the data depicted in Figure 3. It corresponds to phenotypic (DTH) and DL information of genotype 56 observed in 50 environments out of 51 (1-46, and 48-51 but not in env 47). The goal in this example is to estimate DTH for this genotype in the unobserved environment (env 47: Tsukubamirai 2016 Late). For each environment, the planting date was considered as day 1 (*x-axis*) then the daily DL (*y-axis*) values were plotted (pink lines) from day 1 (planting) until genotype 56 reached DTH (blue dots). In Figure 3, a pattern relating DTH on DL at the occurrence day can be identified. This gradient seems not to be strongly affected by the initial DL values at the different planting dates across environments. However, it appears to be linked with the initial DL’s changing rate at the planting day. Those environments where DL’s changing rates decreasing at the planting time (late cultivars) showed shortest DTH values and *vice-versa*.

**3. Figure 3.**
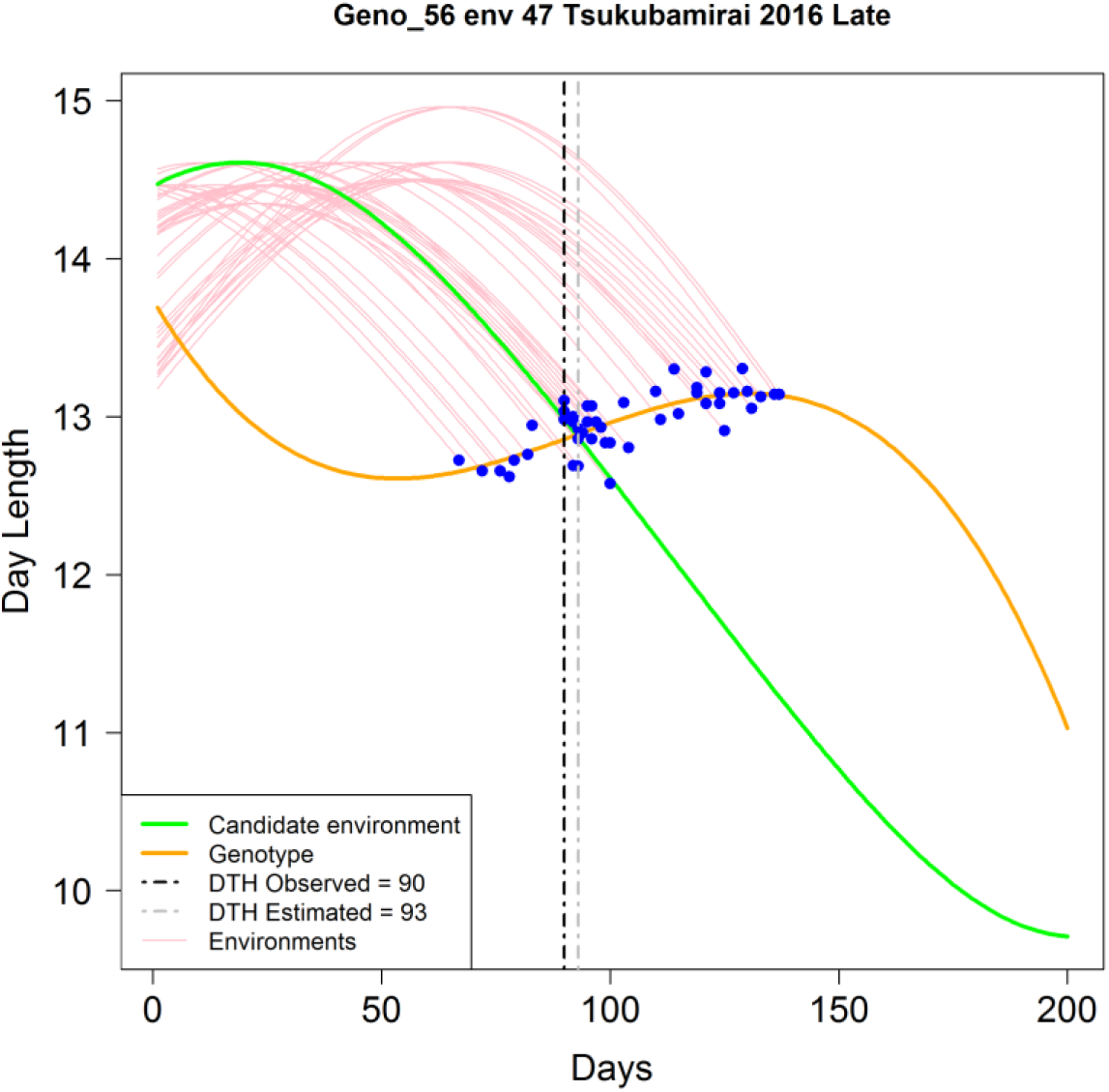
Graphical representation of the C method for estimating days to heading (DTH, *x-axis*) of genotype 56 in environment 47 (env 47 Tsukubamirai 2016 Late). Pink curves represent daily day length in hours (DL, *y-axis*) values from planting day (day 1) until heading time for genotype 56 in 50 environments (1-46, and 48-51). The blue dots indicate the corresponding DL values when genotype 56 reached heading time. The green curve (C_2_) represents the target environment (Environment 47) where no genotypes have been observed yet. The orange curve (C_1_) represents the fitted line for genotype 56 using a third degree polynomial equation relating DL at the occurrence time (DTH across environments). The vertical dotted gray line shows the intersection (93 days) between the green and the orange curves (C_1_, C_2_). This value is the estimated DTH for genotype 56 in the unobserved environment “env 47 Tsukubamirai 2016 Late”. The vertical dotted black line corresponds to the actual DTH (90 days) value for env 47.

For CV00 scheme, the proposed method requires: (*i*) first the estimation of DTH for a set of tested (training) genotypes (one at a time) yet to be observed in a target/unobserved environment (CV0); (*ii*) then the computing of the expected environmental mean of the unobserved environment using the estimated values derived from (*i*) step; (*iii*) and finally to add to the estimated environmental mean the resulting BLUPs of the untested genotypes derived from conventional GS methods. Further details of these steps are given below.

### C method

The DTH estimation of a tested genotype in an unobserved environment (CV0) requires phenotypic and DL information across several environments. The first step considers the characterization of DL at the heading time of occurrence across environments. For this, a third degree polynomial equation is used for fitting DL as function of DTH (orange curve in Figure 3) as follows:

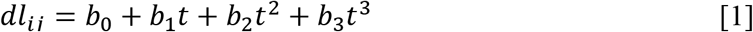

where *dl_ij_* represents the observed DL of the *i*^th^ genotype in the *j*^h^ environment; *t* is the time independent variable; *b*_0_, *b*_1_, *b*_2_ and *b*_3_ are the model coefficients; and *e_ij_* is the error term and it is assumed to be IID *N*(0, *σ*^2^). This model is referred as C_1_.

Similarly, the daily DL information of the target environment (green curve) is modeled using a third degree polynomial equation:

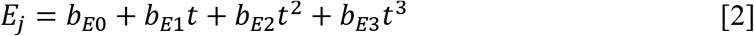

with *b*_0*E*_, *b*_1*E*_, *b*_2*E*_ and *b*_3*E*_ acting as the correspondent model coefficients. The resulting fitted model is referred as C_2_. The solution to this system of two equations (C_1_, C_2_) will produce an estimate of DTH of the genotype in the target environment. In order to find the intersection of these two curves numerical evaluation was implemented. In this example, the solution was found to be 93 days to heading (Figure 3; vertical dotted gray line) while the observed DTH was of 93 days.

### CB Method

Since the ultimate goal is to predict DTH for untested genotypes in unobserved environments, the first step is to accurately estimate the expected DTH mean of the target environment. Then, the final predicted value is composed of the expected DTH mean of the target environment plus the genomic estimated breeding value (*ĝ_t_*) of the untested genotype obtained via conventional GS implementation. For estimating the DTH mean, the C method was first implemented for predicting DTH on the target environment. Here, only the phenotypic and DL information of those genotypes in training set was considered and the predicted values were averaged.

Figure 4 shows the process for predicting untested genotypes in unobserved environments using a toy example consisting of five genotypes (G_1_, G_2_, G_3_, G_4_, and G_5_) and four environments (E_1_, E_2_, E_3_, and E_4_). In this example we describe the procedure for predicting DTH of the untested genotype G_5_ in the unobserved environment E_4_. The top panel of Figure 4 contains the observed values of genotypes 1-4 in environments 1-3; the middle panel shows the predicted values presumably obtained with the C method for genotypes 1-4 (104, 105, 110 and 114) in the unobserved environment 4, also the predicted E-mean (108.3) was computed using as the mean of the predicted DTH values; and the bottom panel shows the predicted DTH for genotype G_5_ in environment 4 (106). This value (106 = 108.3-2.3) was obtained as sum of the Emean and the BLUP value (2.3) obtained for genotype G5 using the conventional GS implementation.

**4. Figure 4.**
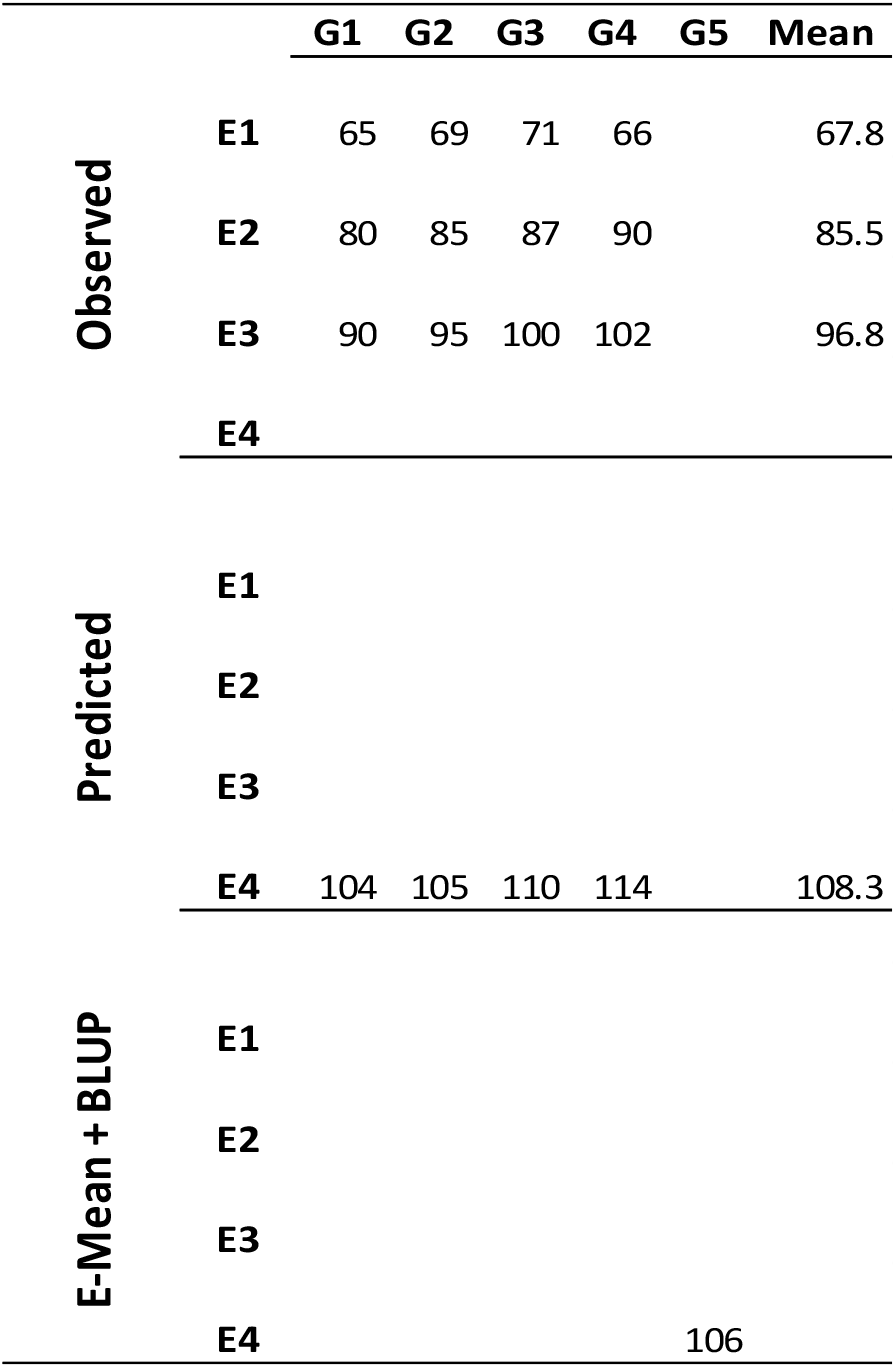
Representation of the procedure for predicting DTH for the untested genotype G_5_ in the unobserved environment E4 using information of four genotypes (G_1_, G_2_, G_3_ and G_4_) observed in three environments (E_1_, E_2_ and E_3_). The top panel contains the observed values for genotypes (G_1_, G_2_, G_3_ and G_4_) in environments (E_1_, E_2_, and E_3_). The middle panel contains the predicted values that presumably were obtained with the C method for these genotypes in the unobserved environment E_4_. Also the E-mean (108.3) of E_4_ is computed as the mean of the predicted values for genotypes G_1_, G_2_, G_3_ and G_4_. The bottom panel contains the predicted DTH value (106) of genotype G5 in environment E4 as the sum of the E-mean (108.3) computed in the previous step and the BLUP (2.3) obtained with the conventional GS implementation.

## Software

The C and CB methods as well all the statistical analyses were performed using the R-software R Core Team (2018)^28^. The conventional GS model described was fitted using the Bayesian Generalized Linear Regression (BGLR) R-package^29, 30^.

## Data availability

Genome data can be accessed from DDBJ under submission DRA008071.

## Acknowledgements

We are grateful to Dr. Vikas Belamkar for their valuable feedback and intellectual contribution during the elaboration of this manuscript.

## Author Contributions

D.J. and H.I. conceptualized the study; H.K.K., S.Y., H.N., and M.Y. performed phenotyping; H.K.K. and C.T. generated and processed molecular marker, environmental and day length data sets; D.J., C.T., R.P. and H.I. analyzed the data; D.J., H.I., R.P., C.T., H.K.K and J.Y. prepared the draft M.S. and all authors contributed to finalize; all authors have read and approved the M.S.

